# Single-cell precision nanotechnology *in vivo*

**DOI:** 10.1101/2023.07.24.550304

**Authors:** Muge Molbay, Benjamin Kick, Shan Zhao, Mihail Ivilinov Todorov, Tzu-Lun Ohn, Stefan Roth, Alba Simats, Vikramjeet Singh, Igor Khalin, Chenchen Pan, Harsharan Singh Bhatia, Farida Hellal, Reinhard Zeidler, Arthur Liesz, Nikolaus Plesnila, Hendrik Dietz, Ali Erturk

**Author notes:** These authors contributed equally to this work (Muge Molbay, Benjamin Kick, Shan Zhao).

## Abstract

Targeting nanoparticle therapeutics with cellular accuracy in whole organisms could open breakthrough opportunities in precision medicine. However, evaluating and fine-tuning the biodistribution of such systems in the whole organism at the cellular level remains a major obstacle. Here, we constructed targetable DNA origami, and analyzed biodistribution in transparent mice, in addition to studying tolerability, clearance kinetics, and immune response parameters. Untargeted DNA origami primarily accumulated in the spleen and the liver, while an immune cell-targeting variant successfully attached to immune cells throughout the body. A cancer cell-targeting mimetic co-localized on solid-tumor metastasis in the liver and the lung. These findings indicate that DNA origami can be directed in vivo, providing an important proof-of-concept and highlights the potential of high-resolution tissue-clearing imaging technologies in their development.

**Graphical Abstract:** 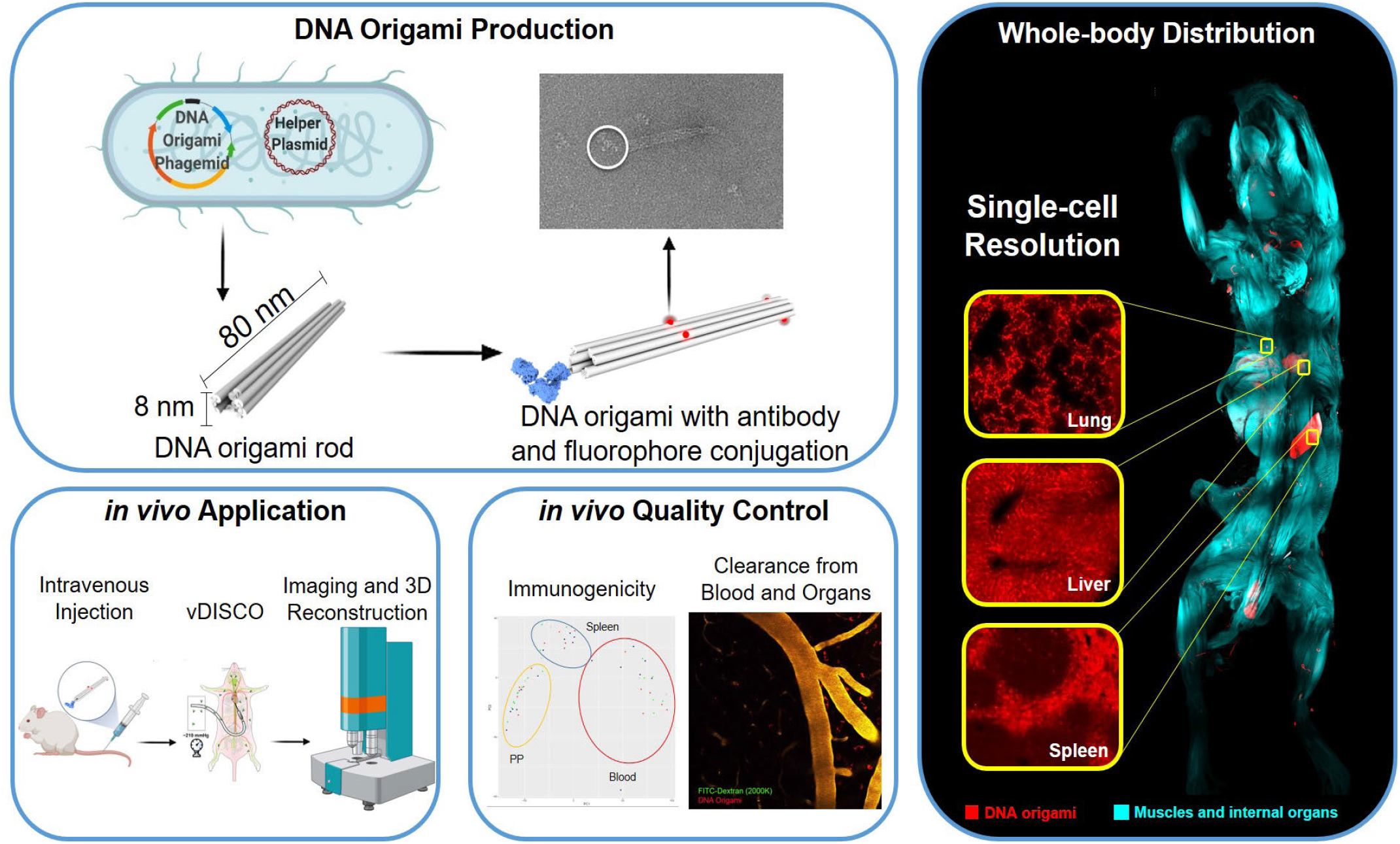

**Highlights:**

1. This study demonstrates the potential of DNA origami-based drug delivery systems as versatile tool for for targeted delivery, which could be used to treat a range of diseases with applications.
2. The immune compatibility, half-life, targeting efficiency, and the biodistribution evaluation of DNA origami indicate its potential for systemic drug delivery.
3. Our approach enables the assessment of biodistribution of nanoparticles in the intact body with a sensitivity to the single-cell level, highlighting the high-resolution tissue clearing technoloies in revealing DNA origami’s feasibility for drug targeting.

## INTRODUCTION

Precision medicine has been a major focus of recent biomedical research, primarily identifying subgroups of patients and targeting the disease-carrying cells (1, 2). Recent advances in multi-omics approaches, organoids, and artificial intelligence (AI) now enable the identification of disease heterogeneity within patients with their distinct types of diseased cells (1, 3–5). However, the next level of precision in medicine, i.e., delivering drugs specifically to those subtypes of cells needing specific treatments, has been comparably slow to advance. Diverse solutions, including nanoparticles, have been proposed to undertake cell-level targeting to accomplish this task. However, developing precision delivery systems were hampered by comparably coarse resolution biodistribution data obtained by imaging methods such as Positron Emission Tomography (PET) and bioluminescence at the whole mouse body level, whose resolution is far from the needed cell level accuracy.

Programmable self-assembly with DNA origami (6–9) enables the bottom-up assembly of designer nanoparticles with emerging applications in various fields (9–12). DNA origami particles show fundamental characteristics that are highly attractive for precision drug delivery systems: biocompatibility, control over shape and dimensions of the designed object, the site-directed functionalization with organic and inorganic compounds almost throughout the entire structures. Moreover, this offers the possibility to create objects that reconfigure in response to molecular stimuli. Many innovative concepts for new types of vaccines (13), antivirals (14, 15), and stimuli-responsive precision drugs (16, 17) have been envisioned using DNA origami. However, evaluating the applicability of these concepts in actual organisms remains in its infancy. It is also only recently that structurally stabile DNA origami for application in physiological fluids (18, 19) and for producing DNA origami at biomedical quantities (20) and free from bacterial endotoxins have been discovered.

In parallel, developments in tissue clearing and lightsheet microscopy now allow imaging at cellular details in whole organs and rodent bodies, and even human embryos and organs (21–25). In particular, vDISCO technology renders whole mouse bodies transparent by combining decalcification, decolorization, and organic solvent-based tissue clearing. Furthermore, it can enhance the signal of the endogenously expressed fluorescent proteins by two orders of magnitude to enable visualizing cellular details throughout the intact transparent mouse bodies assessed (26,27).

Here, we utilized whole mouse transparency and panoptic imaging to design and finetune targeting of DNA origami drug mimetics at a single cell level in the whole mouse body. To this end, we generated bioreactor-produced DNA origami rods (20) as a model. for mass-producible future DNA therapeutics, including removing E. coli lipopolysaccharide endotoxins for immune tolerability covalent conjugation with fluorescent labels and targeting antibodies, and coating polymers for enhanced *in vivo* stability. After preparing the DNA origami drug mimetics, we injected them in mice and assessed their clearance kinetics and biodistribution at the cellular level using vDISCO whole-body transparency (26,27). We also evaluated the pharmacokinetics and immune response over time to further substantiate our approach to develop DNA origami drug mimetics that are safe and functional at a single-cell level *in vivo*.

## RESULTS

### DNA origami drug mimetics for in vivo tracing

We employed the methods of multi-layer DNA origami (2) to build three variants of a ten helix bundle in honeycomb lattice packing based on a single-stranded 2581 bases long DNA scaffold and 21 single-stranded DNA staples with an average length of 115 bases, as previously described (20). The resulting particles were 80 nm long and approximately 8 nm in diameter (Fig. S1A). To detect the DNA origami particles in vivo, we labeled them with multiple Atto550 or Atto647 dyes. We coated the DNA origami with poly(ethylene glycol)–poly(L)Lysine (PEGPLys) block copolymers (18) to enhance structural stability during circulation, which verified first ex vivo in 5% and 80% mouse serum-containing PBS prior to in vivo experiments (Fig. S1A). The particles were further purified from E. coli lipopolysaccharide endotoxins. All administered samples were below 36 EU/ml for a daily dose of 100 μl satisfying the standard residual concentration of endotoxins for in vivo applications (28). As a result, we obtained fluorophore-conjugated, PEG-poly(L)Lysine coated DNA origami rod variant as the base structure, which we refer to as the “non-targeting origami”. We also prepared two variants designed to target distinct cell types (Fig. S1B-E): The “immunecell-targeting origami,” which included one rabbit anti-CX3CR1 IgG antibody to target cells of the immune system (mainly monocytes and macrophages). The anti-CX3CR1 antibody binds to amino acids 175-189 of the receptor for the CX3C chemokine fractalkine in lymphoid and neural tissues (29). The “cancer-targeting origami” included two human carbonic anhydrase (CA) XII-specific IgG antibodies (6A10) to target cancer cells that over-express the carbonic anhydrase (Fig. S1F). In both cases, DNA-conjugated antibodies were attached using single-stranded DNA extensions protruding from the origami rod. The cancer-targeting variant was labeled with Atto550 instead of Atto647 to avoid spectral overlap with cancer metastases signal at 647 nm.

### Bio-distribution of origami rods in whole mouse body

So far, there have been no reports to visualize targeting of nanotechnology-based drug carriers in the whole mouse body at the cellular level, preventing the assessment and fine-tuning of such systems for cell level targeting. To analyze the DNA origami distribution in vivo at the cellular level (26), we injected “non-targeting origami” and “immune-cell-targeting origami” through the femoral vein in C57BL6 mice. We injected “cancer-targeting origami” in mice carrying metastatic mammary carcinoma cells (MDA-MB-231). As the tumor cells were transfected to express both mCherry and firefly luciferase (27), we could follow cancer progression by bioluminescence before the cellular level observation of mCherry signal using vDISCO (Fig. S2). After allowing for circulation of the injected DNA origami (20 min for non-targeting and immune-cell targeting, 60 min for cancer-targeting), the mouse bodies were collected vDISCO transparency, stained with propidium iodide (PI) to label the cell nuclei, and imaged using light-sheet microscopy (26). For the cancer model mice, we boosted the signal of mCherry-expressing MDA-MB-231 cancer cells to image them along with DNA origami instead of PI labeling using light-sheet microscopy. The non-targeting origami was primarily distributed in the liver (Fig. 1A, arrow), while less present in other organs such as lung, heart, kidney, and lymph nodes, suggesting a non-specific binding during hepatic filtration (Fig. 1A, arrowheads). The signal from the immune-cell-targeting origami appeared different already at the whole mouse level. It was identified predominantly in immunological organs such as lymph nodes at various body sites (Fig. 1B, arrowheads) and the spleen (Fig. 1B, arrows, Movie S1), suggesting binding to CX3CR1+ monocytes/macrophages. The cancer-targeting origami signal again had a biodistribution distinct from those of non-targeting and immune-cell-targeting variants: the cancer-targeting origami mainly accumulated in the **l**ungs and other regions in the body where cancer metastases were observed (Fig. 1C, arrows).

**Figure 1.**
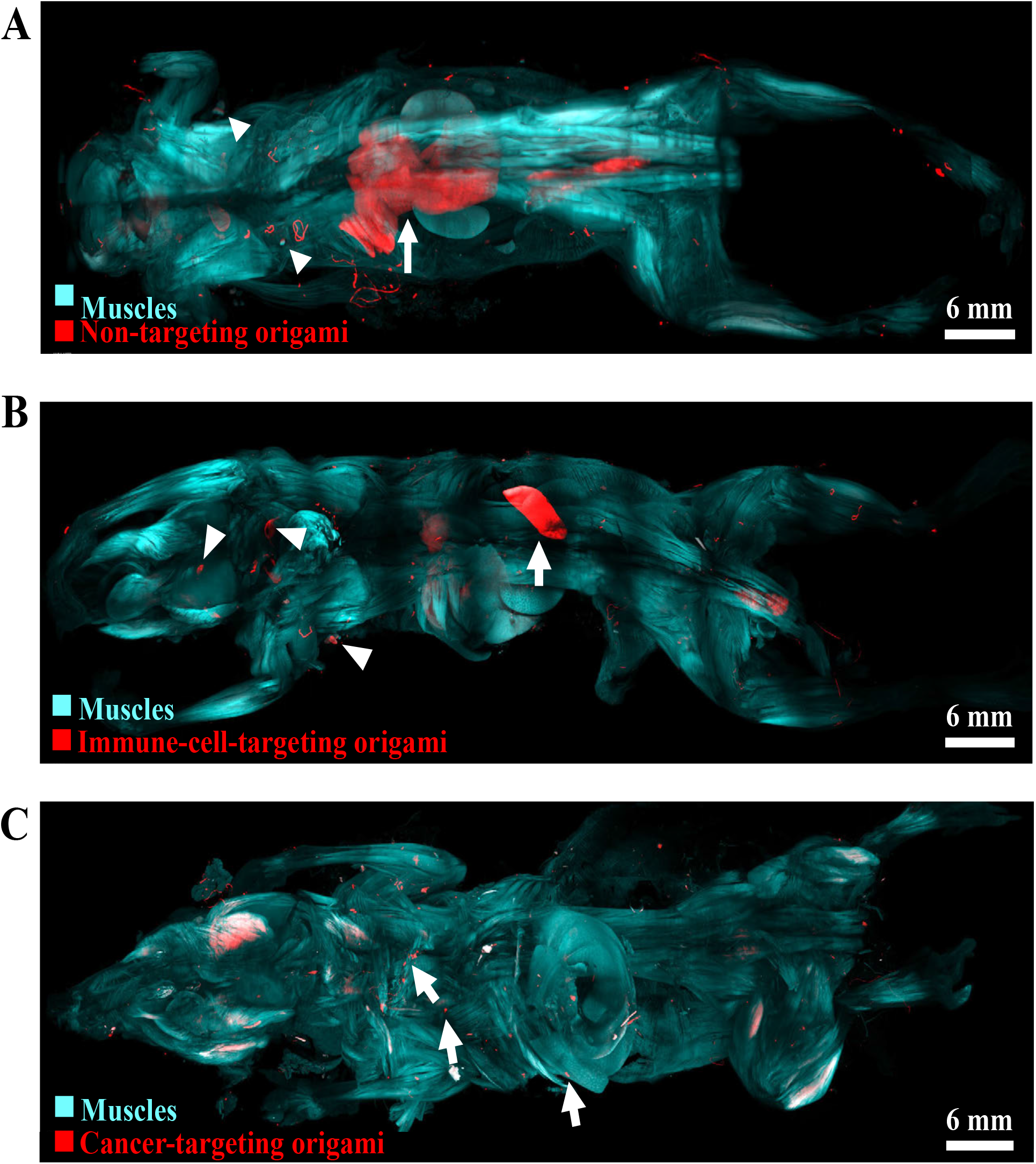
*in vivo* distribution of DNA origami rod-3D reconstruction of DNA. The autofluorescence of muscles scanned at 488 nm is used to highlight the shape of whole mouse body in each example. **a**, Non-targeting origami (basic rod structures) are mainly observed in the liver (arrow), and to less degree in immune organs such as lymph nodes (arrowheads). **b**, Immune-cell-targeting DNA origami, conjugated with CX3CR1 antibody, targets monocytes, macrophages and microglia. In this case, DNA origami signal become stronger in diverse organs including liver and spleen (arrows). **c**, Cancer-targeting DNA origami (red), conjugated with 6A10 antibody, targets cancer micrometastases throughout the mouse body (arrows).

We further compared the origami distribution at the organ level, particularly in the lungs, liver, and spleen, having stronger signals over other regions. As a control, we also imaged animals where only Atto-labeled oligonucleotides (Atto-ON) alone were injected (no DNA origami) and found no detectable signal (Fig. 2A). This suggests that the Atto-oligonucleotides cannot accumulate in the organs and are excreted from the body rapidly. This data also provides evidence for the stability of observed origami-Atto structures and their specificity. The nontargeting DNA origami signal was present in the liver and spleen but almost undetectable in the lungs (Fig. 2B). The immune-cell-targeting origami displayed higher signal intensity in the lungs, liver, and spleen than non-targeting origami indicating increased tissue retention (Fig. 2C). In the liver, non-targeting and immune-cell targeting origami particles appeared within the hepatic sinusoids where the Kupffer cells, the liver resident macrophages, are in contact with endothelial cells (30). In the spleen, the immune-cell-targeting origami primarily accumulated in the red pulp, a large reservoir for monocytes bearing CX3CR1 (31). In contrast, the cancer-cell targeting origami signal was absent from the liver and spleen but present in the lung (Fig. 2D) and in the liver as clusters (Fig. 2E), where most of the metastasis for this cancer line occurs (32). These results demonstrated that the DNA origami particles are indeed directable in vivo to specific organs and tissues, and targeting moiety can control their distribution.

**Figure 2.**
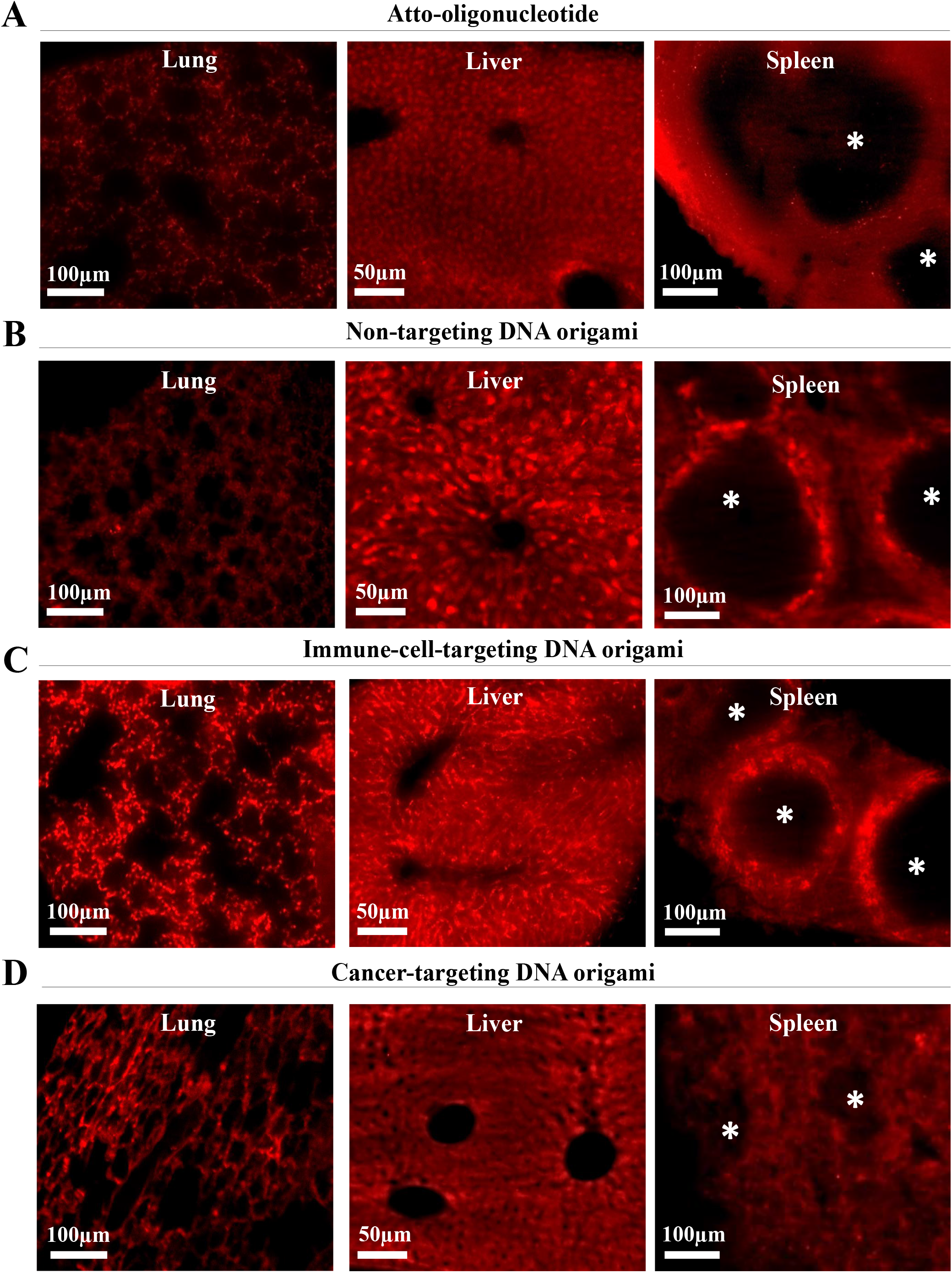
Distribution of DNA origami with different targeting moieties in selected organs. **a**, Atto-oligonucleotide, **b**, non-targeting DNA origami, **c**, immune-celltargeting DNA origami, and **d**, cancer-targeting DNA origami. All organs are imaged at 12x magnification; red signal represents Atto dye with or without DNA origami. The asterisk point the signal free white pulp of spleen.

### Origami distribution at the cellular level

After identifying the different origami constructs at the whole mouse and organ level, we dissected the tissues for a sub-cellular level analysis using confocal microscopy. We found that the non-targeting and immune-cell-targeting DNA origami signals had distinct distributions in the liver tissue. The non-targeting origami localized mainly within the sinusoids of the liver (Fig. 3A). These liver regions are enriched in endothelial cells and Kupffer cells and are responsible for the translocation drugs to hepatocytes, the site of their metabolism (33). The immune-cell-targeting DNA origami distribution in the liver is similar to the naked DNA origami.However, it has higher signal intensity and deeper tissue penetration (Fig. 3b), indicating better retention of the DNA origami when conjugated with immune cell targeting antibodies. The key actor of the similarity in the distribution pattern in two moieties is the Kupffer cells. They are the phagocytic cells of the liver.

**Figure 3.**
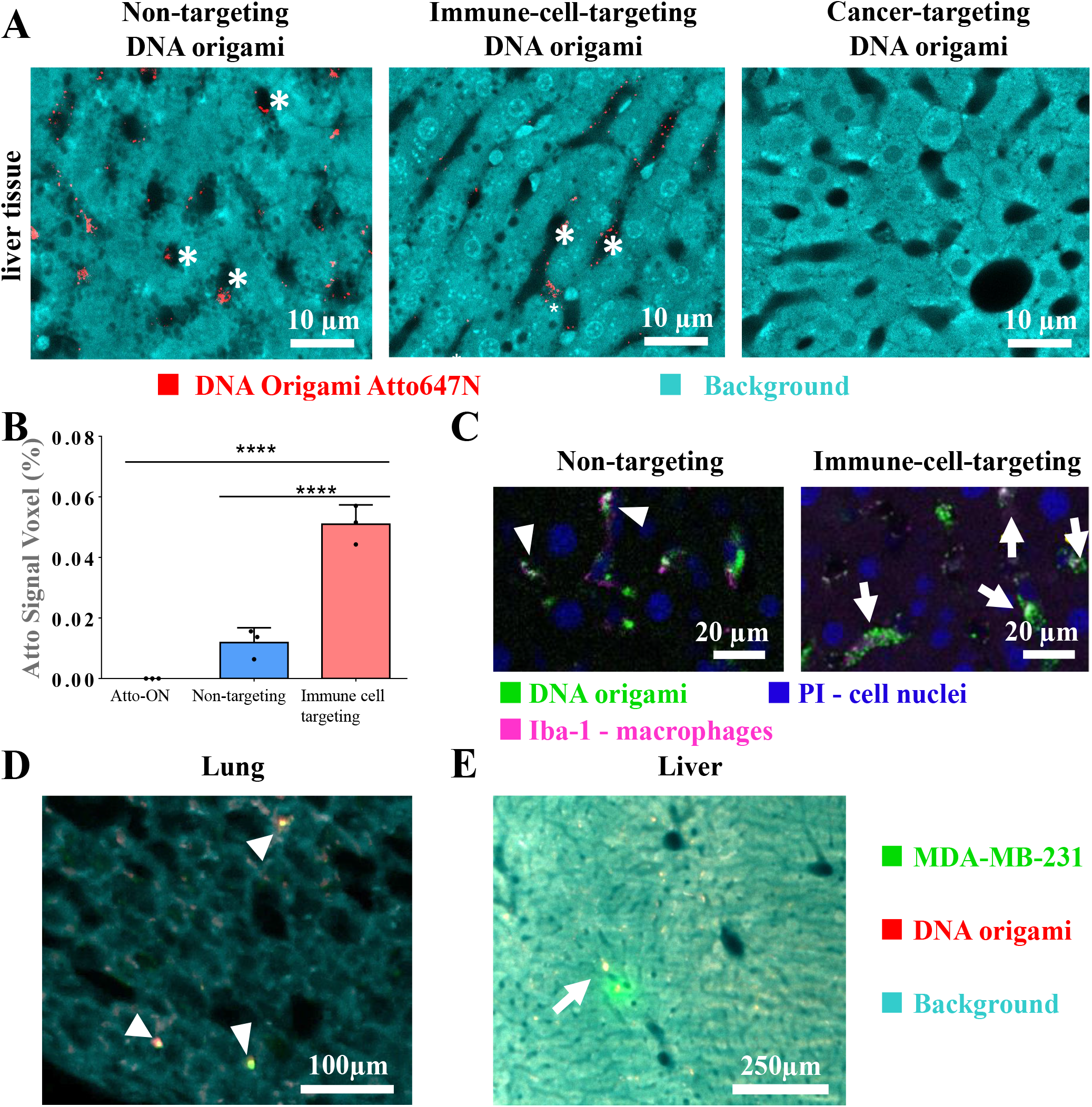
Comparison of the cellular localization of different DNA Origami. **a**, Confocal images of liver from non-targeted, CX3CR1 targeting and 6A10 conjugated origami. Background is blue, PI-labelled nuclei red and DNA origami signal green. The asterisks point the liver sinusoids, which is a type of capillary with a lining of endothelial cells and Kupffer cells (CX3CR1+). **b**, Quantification of signal voxel (%). CX3CR1 targeting DNA origami group has significantly higher signal in liver (Data are presented as mean ± SD; ****:p < 0.0001; one-way ANOVA). **c**, Subcellular addressability of DNA origami for CX3CR1 positive tissue macrophages evaluated by immunohistochemistry in 1 mm sections of livers with macrophage specific Iba-1 antibody in naked DNA origami Atto647 or CX3CR1 targeting origami injected animals. (Blue: PI-labelled nuclei; Magenta: Iba-1 antibody showing macrophages; Green: Atto647 dye). Engulfment of DNA origami CX3CR1 antibody Atto647 PEG-Polylysine with higher efficiency can be observable with co-localization of DNA origami signal and Iba-1 or CD68 labeling (arrows). **d**, The view from the lung and liver **e**, in the cancer mice showing that DNA origami signal is overlapping many regions with cancer metastases (arrow), and there are some non-targeted MDA-MB-231 cells (arrowhead).

Additionally, Kupffer cells are CX3CR1 positive, and they are attached to the walls of sinusoids; therefore, immune cell-targeting DNA origami localizes the same region. To obtain detailed, sub-cellular information on the localization of DNA origami, we also labeled the tissues with antibodies targeting either Iba-1 or CD68, which are markers for monocytes and macrophages, respectively. Iba1 is expressed in the cytoplasm, whereas CD68 in lysosomal and endosomal vesicles (34). The non-targeting origami displayed indiscriminate staining, mainly located within the cells, while the CX3CR1 conjugated DNA origami co-localized with Iba1 (Fig. 3C) and CD68 (Fig. S3) positive cells. Internalization of non-targeting DNA origami suggests random phagocytosis and the immune-cell-targeting origami covering the surface of the IBA1 positive cells, suggesting that the origami decorated the surface membrane of these cells at the CX3CR1 protein location.

We also investigated cancer-targeting origami conjugated with 6A10-antibody origami at the subcellular level. We found many micrometastases in the lung, liver, and kidney (Fig S2B). High magnification imaging of lungs revealed punctate signals from the cancer-targeting origami colocalized with some but not all the cancer cells present in the tissue (Fig. 3D). Similarly, metastases from the liver co-localized with the cancer-cell targeting origami particles (Fig. 3E). Our data show that DNA origami is targetable to intended cells at the cellular level.

### Immune response, physiological and pharmacokinetic parameters of DNA origami used at the treatment doses

A significant challenge for the in vivo use of nanocarriers is the induction of adverse effects such as excessive immune response (36). As we used therapeutic quantities of DNA origami, we assessed the level of immune response 4 and 24 hours after DNA-origami injection. To this end, we performed an immunophenotypic analysis via flow cytometry and focused on early-activated T cells, characterized by CD45+ CD3+ CD69+, along with the CD45+ CD11b+ monocytes. DNA origami without any particular action to remove residual E. coli endotoxins induced a substantial immune response: T cells and monocytes numbers were highly increased in blood and spleen. In contrast, the purified DNA origami, which had an endotoxins concentration of 0.5 EU/ml, did not induce a significant increase of active T cell or monocyte numbers compared to the mice treated with buffer controls (Fig. S4). We further assessed the physiological effects of orally administered doses of DNA origami in CD1 animals. The body weight, body temperature, and basic sensorimotor abilities did not change during the observation time of seven days (Fig. 4A, B, and C). These data suggest that DNA origami does not affect the general physiological parameters of the animals. Thus, the non-targeting E.coli-endotoxin purified DNA origami was neither toxic nor did it trigger a significant immune response up to 7 days after administration. For completeness, in another group of animals undergoing the same treatment, we collected the peripheral blood and spleen at 24 hours and at seven days to assess the long-term immune response upon orally administrated DNA origami. We evaluated immunogenicity via flow cytometry focusing on the previously mentioned activated T cells (CD45+CD3+CD69+) and monocytic cells (CD45+CD11b+). The percentage of MHCII+ cells acquired monocytic cell activation. We did not observe any alterations in the total T cell count or activated T cells and monocytes neither at 24 hours nor at seven days after gavage in the blood (Fig. 4D) or spleen (Fig. 4E).

**Figure 4.**
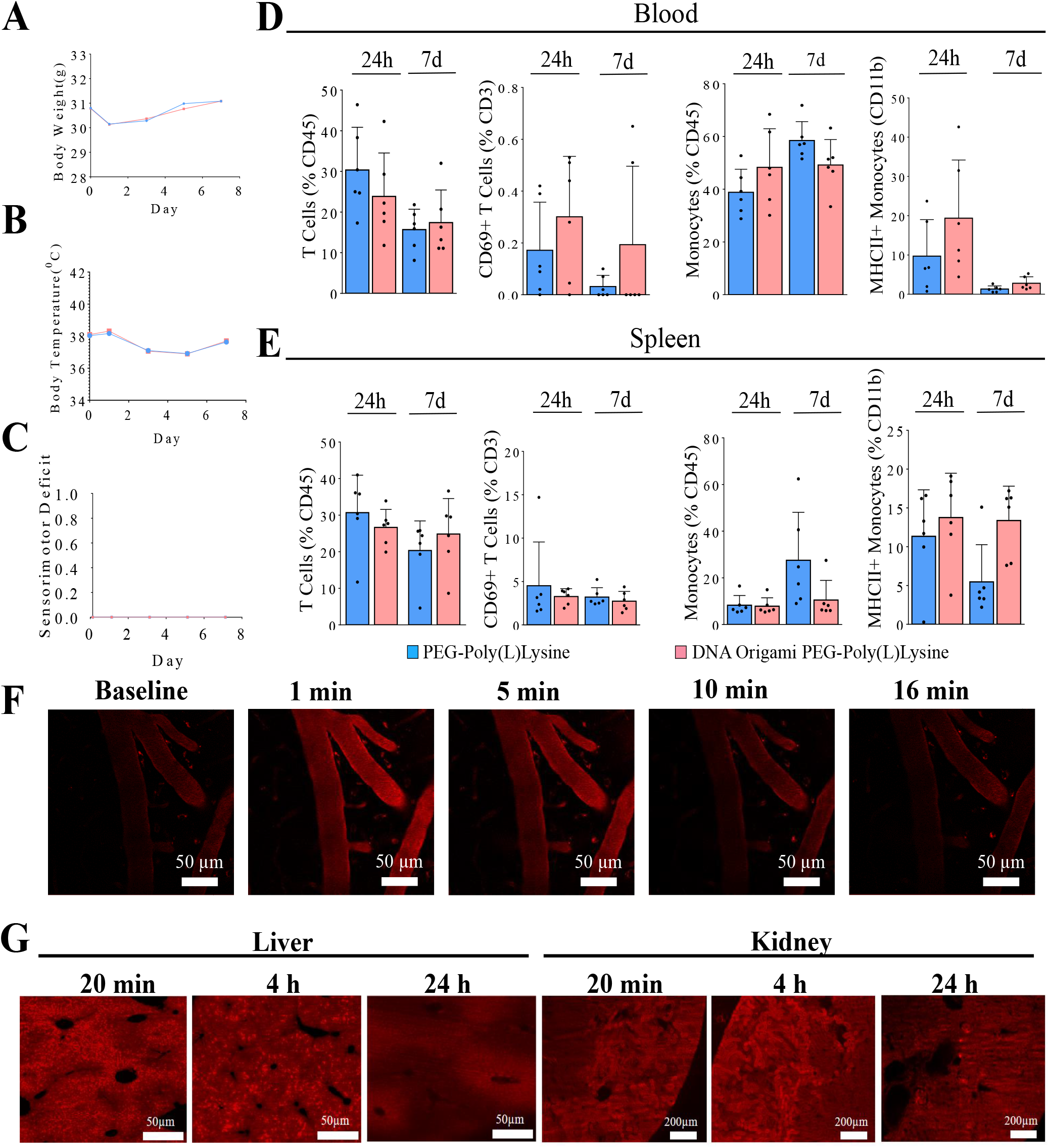
Immune parameters and in vivo clearance. Effects of DNA origami and PEG-Poly(L)Lysine control, administered by gavage on **a**, body weight, **b**, body temperature and **c**, behavioral score (on day 0, 1, 3, 5 and 7). Comparison of monocytes and T cell populations and activation in **d**, blood and **e**, spleen after DNA origami administration **f**, Blood clearance of naked DNA origami by intravital 2-photon imaging of the pial vessels after intravenous injection of non-targeting DNA origami (red) for 70 minutes. Clearing of DNA origami from the **g**, liver and **h**, kidney, 4 hours and 24 hours after femoral vein injection and 20 minutes circulation.

To elucidate the pharmacokinetic profile of the DNA origami particles, we used in vivo 2-photon imaging of the pial vessels and assessed the blood clearance of the non-targeting variant. To this end, we implanted an acute cranial glass window over the cerebral cortex of C57BL6 mice and inserted a femoral arterial catheter for the injections (37). The animals were placed under a 2-photon microscope and received FITC Dextran (2000 KDa) injection to label the cerebral vessels to acquire a baseline signal. After the injection of non-targeting origami, the fluorescent signal was recorded for the next 70 minutes (Fig S5, Movie S2). The origami signal immediately increased upon injection and returned to baseline level within 16 minutes (Fig. 4F). The clearance timescale suggests that the DNA origami particles have low binding to circulating plasma proteins of mouse blood serum and readily transition from the blood to the vessel walls and solid tissues, as we observed above by whole mouse imaging.

To test the kinetics of clearance of non-targeting origami from internal organs, we injected particles through the femoral vein of the mice and let them circulate for 20 minutes, 4 hours, and 24 hours before sacrifice. After 20 minutes of circulation, the non-targeting origami signal was already visible in the liver tissue (Fig. 4G). It appeared as a speckled pattern radially distributed from the blood vessel lumens to the tissue. The signal intensity decreased at 4 hours. After 24 hours, no signal above the background was visible. We noted that no appreciable origami signal was observed in the kidney at any time, indicating a continuous excretion of the particles without accumulation in the kidney (Fig. 4H). These data suggest that DNA origami amounts peak in the solid tissues and organs within the first few hours of injection and gets eliminated throughout approximately 24 hours. This kinetic sets a reasonable time window for drugs to reach their targets before clearance of their biodegradable origami carriers is eliminated.

## DISCUSSION

While advanced nanotechnologies promise to overcome hurdles around delivering treatments to pre-specified cells, so far, assessing the biodistribution of nanoparticles across whole organs or even the organism relied on imaging methods such as PET, bioluminescence, and MRI providing resolutions far away from the cellular level. Whereas cell-level resolution in direct imaging would benefit, maybe is even critical, for evaluating and fine-tuning nanoscale nanoparticles for precision medicine to maximize on-target activity and minimize off-target effects. Conventionally, tissue histology analyzes selected tissue sections with higher resolution. However, such an approach ignores ∼99% of animals’ tissues bearing the danger of introducing substantial bias, e.g., missing to identify all tissues being targeted and affected in the whole organism. Here, we present the first study outlining a whole-organism approach to assess and fine-tune cellular level targeting nanocarriers using whole mouse body transparency and DNA origami.

DNA origami nanoparticles are modular and biodegradable nanostructures (5,48) that offer attractive molecular engineering opportunities to help overcome the limitations of conventional drugs for targeting and improving disease therapy specificity. Here we addressed the safety, biodistribution and clearance, and the targeting of engineered DNA origami drug mimetics (“chassis”) in whole mouse bodies at cell level resolution by taking advantage of recent advanced whole-body transparency and imaging technologies (6,7). We did not observe any adverse effect in the tissues or in the behavior of mice treated via intravenous route or gavage with the DNA origami with or without antibody attached, suggesting biocompatibility and safety for medical usage in principle. However, independent from the targeting moiety, DNA origami seems to be unable to penetrate through the blood-brain barrier (BBB) as we have not noted any significant origami signal in the brain parenchyma. The current effort focuses on combining DNA origami with different coating materials to hijack BBB. Additionally, taking advantage of multi-omics studies and using resulting specific molecular signatures to enhance the targeting to a more focused group of cells can provide more precise and personalized treatment options. Furthermore, thanks to the very high cargo capacity and programmability of DNA origami, a combination of targeted DNA origami with multiple functional molecules as drugs, inhibitors, etc., and monitoring in vivo efficiency of these complex particles will widen the horizons of in vivo nanoparticle applications.

In conclusion, we present the first methodological whole-organism approach for the development of nanocarriers that are controllable at the cell level. While we tested our approach exemplarily with DNA-based nanoparticles, the pipeline applies in principle to any other nanoparticles. Future studies can use this approach to develop tailored treatments where the cell-level accuracy is highly critical, including treatment of cancer metastasis and somatic gene editing of user-specified cells.

## MATERIALS AND METHODS

### DNA origami production

The biotechnological produced nanorod was prepared as recently described (49). In brief, the staple strands for the nanoobject are arranged as a pseudogene, interleaved by self-excising DNAzyme cassettes. The pseudogene is cloned into a phagemid backbone, which functioned as a scaffold strand for the nanoobject. The fully cloned phagemid was introduced into E. coli JM109 cells by transformation. Using helper phage rescue, the phage particles containing the phagemid single-stranded DNA are produced in a stirred-tank bioreactor, followed by phage and DNA purification to yield pure circular ssDNA. As a prerequisite for the in vivo experiments, the produced ssDNA was endotoxin purified as described by Hahn50, using Triton X (114). By adding zinc cations, the self-excising DNAzyme gets active and yields scaffold and staple strands for the nanorod. After a further DNA purification step with Ethanol the nanorod was assembled in a buffer containing 5 mM Tris, 5 mM NaCl, 1 mM EDTA, and 10 mM MgCl2 at a DNA concentration of 50 nM (15 min at 65°C, then 60-45 °C at 1 h/°C. To assemble fluorescent-labeled DNA nanoobjects for the vDISCO protocol, we purchased chemically synthesized single-stranded staple strands with Atto 550 or Atto 647N modification from IDT or Biomers. The folding reactions were set up as 100 μl or 2 mL reaction solutions, dependent on the amount necessary for the in-vivo experiments.

### Preparation of IgG-ssDNA conjugates

Oligonucleotides modified with 3’ thiol modification were purchased HPLC purified and dried by Biomers (Germany). The oligos were dissolved in PBS (100 mM NaPi, 150 mM NaCl, pH 7.2) with 5 mM TCEP and incubated for 1h at RT. After three rounds of filter purification (10k Amicon Ultra 0-5 mL centrifugal filter), 10 nmol of the reduced thiol oligo was mixed with 10 equivalents of Sulfo-SMCC (sulfosuccinimidyl 4-(N-maleimidomethyl)cyclohexane-1-carboxylate, Thermo, dissolved in ddH2O) for 15 min. After three rounds of filter purification (10k Amicon Ultra 0-5 mL centrifugal filter), including buffer change to PBS (pH 8), 100 μg of antibody in PBS (pH 8) was added. The reaction was incubated for 4 h at 4°C at least. The conjugate was subsequently purified by ion-exchange chromatography (proFIRE, Dynamic Biosensors) using a NaCl gradient of 150-1000mM in PBS (pH 7.2). The purity of the oligo-antibody conjugates was analyzed by SDS-PAGE and agarose gel.

### Assembly of antibody functionalized nanorod

The assembled nanorod was purified with two rounds of PEG precipitation to remove excess staple strands as described by Evi Stahl, with slightly higher PEG 8000 (8.25 % w/v) and NaCl (275 mM) concentrations51. The nanorod was equipped with up to 4 handle ssDNA strands with complementary sequences to the IgG-ssDNA conjugate. The hybridization of IgG-ssDNA conjugate to the nanorod was done in buffer containing 5 mM Tris, 5 mM NaCl, 1 mM EDTA, and 10 mM MgCl2 at pH 8 overnight at 30°C (see Supplement Figure 1 B). To concentrate the fully functionalized nanorod, we use another round of PEG purification, followed by dissolving the nanorod in Tris buffer (5 mM Tris, 5 mM NaCl, 1 mM EDTA, and 5mM MgCl2) at a high concentration of 4 μM. We added PEG-polylysine (methoxy-poly(ethylene glycol)-block-poly(l-lysine hydrochloride, 10 lysine repeating units, 5 kDa MW PEG, Alamanda polymers) at an N:P ratio of 1:1 to stabilize the nanorod against low salt conditions and nuclease activity in vivo. For the in vivo experiments, the nanorod was diluted to 0.5 μM or 2 μM using sterile PBS to reduce the magnesium concentration to at least 2 mM or less. Finally, the endotoxin concentration of the fully equipped nanorod was determined using Endosafe Nexgen-PTS (Charles River) to fulfill the FDA requirement of 36 EU/ml for a 100 μl interveinal injection per day (37).

### Animals

We used the following animals in the study: mixed-gender CD-1 IGS mice (Charles River, stain code: 022), C57BL/6J (Jackson Laboratory strain code: 000664). Female NSG (NOD/SCID/IL2 receptor gamma chain knockout) mice were obtained from Jackson Laboratory. The animals were housed under a 12/12 hours light/dark cycle. The animal experiments were conducted according to institutional guidelines: Klinikum der Universität München / Ludwig Maximilian University of Munich and after approval of the Ethical Review Board of the Government of Upper Bavaria (Regierung von Oberbayern, Munich, Germany) and the Animal Experiments Council under the Danish Ministry of Environment and Food (2015-15-0201-00535) and following the European directive 2010/63/EU for animal research. All data are reported according to the ARRIVE criteria. Sample sizes were chosen based on prior experience with similar models. Sample sizes are specified in figure legends. Within each strain, animals were randomly selected. Animals that resulted negative for the expression of fluorescent proteins by genotyping were excluded from the study.

### Immunotoxicity assessments

Mixed-gender CD-1 mice (n=5) received tail-vein injections of naked DNA origami (2 μM x 100 μl for each mouse, 13,68 mg/kg DNA origami per animal with and without purification). After 4h and 24h, the blood and spleen were collected. The mice (n=5) received gavage injections of DNA origami 2μM x 400μl for each mouse (50,725mg/kg per animal); after 4h and 24h, animals were sacrificed, the blood samples and spleen were collected for flow cytometry analysis of activated immune cells. On day 1, 3, 5 and 7, body weight, body temperature measurements, and basic neurological assessments were done to better examine the effects of DNA origami injection on the body. Flow cytometry was performed in the same way for both experiments. After blood collection. erythrocytes were lysed using isotonic ammonium chloride buffer. Immediately following a cardiac puncture, mice were perfused with normal saline for dissection of the spleen. Spleens were transferred to tubes containing Hank’s balanced salt solution (HBSS), homogenized, and filtered through 40 μm cell strainers to obtain single-cell suspensions. Homogenized spleens were subjected to erythrolysis using isotonic ammonium chloride buffer. Anti-mouse antibodies were used for surface marker staining of CD45+ leukocytes, CD45+CD11b+ monocytes (+ MHCII+ expression), CD3+ T cells and CD3+CD69+ activated T cells. Fc blocking (Anti CD16/CD32, Invitrogen, US) was performed on all samples prior to extracellular antibody staining. All stains were performed according to the manufacturer’s protocols. Flow cytometric data was acquired using a BD FACSverse flow cytometer (BD Biosciences, Germany) and analyzed using FlowJo software (Treestar, US).

### Non-targeting DNA origami half-life in blood

For multiphoton imaging, we used an upright Zeiss LSM710 confocal microscope equipped with a Ti:Sa laser (Chameleon Vision II) from Coherent (Glasgow, Scotland) and two external photomultiplier detectors for red and green fluorescence. All animal experiments were conducted in accordance with institutional guidelines and approved by the Government of Upper Bavaria. 8-week old C56/Bl6N mice obtained from Charles River Laboratories (Kisslegg, Germany) were anesthetized intraperitoneally (ip) with a combination of medetomidine (0.5 mg/kg), fentanyl 11 (0.05 mg/kg), and midazolam (5mg/kg) (MMF). Throughout the experiment, body temperature was monitored and maintained by a rectal probe attached to a feedback controlled heating pad. A catheter was placed in the femoral artery to administer the fluorescent dye or NPs. A rectangular 4×4mm cranial window was drilled over the right frontoparietal cortex under continuous cooling with saline, as it was described52. In order to identify the brain vessels and obtain a baseline image, the mouse was injected with FITC dextran 3μL/g. Afterward, the animal was placed on the multiphoton microscope adapted for intravital imaging of small animals. Non-targeting DNA origami, conjugated with Atto-550, was injected at a concentration of 1μM and a dose of 120 μL. The scanning was performed with time series, 80 μm depth, laser power 10%, 800 nm, GAASP detector with LP˂570 nm filter, and master gain 600 for the FITC channel and LP˃570 nm for the NPs channel with master gain 600. In total, imaging time was not exceed 70 min. The fluorescence analysis was performed using FIJI software.

### Biodistribution and clearance of DNA origami

Catheters were made from polyethylene tubing (inner diameter, 0.28 mm; outer diameter, 0.61 mm); were heated and pulled to obtain a cone-shaped tip. Catheter tips were inspected under a stereomicroscope (magnification, 31.5×; SZX 10, Olympus Schweiz, Volketswil, Switzerland). The tips of the catheters used had an outer diameter of 110 μm ± 30 μm (n = 44) and were angled by using a scalpel. Capillaries were flushed then with sterile 0.9% NaCl. CX3CR1GFP-/+ mice were anesthetized by using 5% isoflurane (Forene, Abbott, Baar, Switzerland) in oxygen (300 mL/min). Anesthetized mice were laid on their backs on a heating pad (Horn, Gottmadingen, Germany), and the body temperature was maintained at 37 °C. During surgery, anesthesia was maintained by using an inspiratory isoflurane concentration of 2%. Surgery was performed under a stereomicroscope (magnification, 31.5×; SZX 10, Olympus). The left femoral artery, vein, and nerve were exposed through a skin incision of 3 to 4 mm parallel and inferior to the inguinal ligament. The femoral vein was separated from surrounding tissues; a small hole was created in the vein approximately 2 to 4 mm distally the intersection with the inguinal ligament. The capillary was replaced with polyethylene tubing by advancing the thinned tip into the artery. Through this capillary 100μl was injected with the following solutions; MgCl2 buffer, Atto647-PEG, DNA Origami-Atto647-PEG (0.5 μM), and goat anti-mouse CX3CR1 antibody conjugated DNA origami (0.5 μM). Animals were sacrificed after 20 minutes, 4 hours, or 24 after injection.

### Administration of cancer targeting DNA origami

Cancer cell injection and bioluminescence measurement of tumor size were done following the DeepMACT protocol (53). MDA-MB-231 breast cancer cells transduced with a lentivirus expressing mCherry and enhanced-Firefly luciferase were counted, filtered through a 100 μm filter, and resuspended in ice cold DPBS in the concentration of 1×106 cells/mL. 100μL of this cell suspension was injected to immune deficient NSG (NOD/SCID/IL2 receptor gamma chain knockout) mice via transcardiac route. Tumor growth was monitored just before DNA origami injection by bioluminescence measurement (photons/second) of the whole body using an IVIS Lumina II Imaging System (Caliper Life Sciences). Briefly, mice were anesthetized with isoflurane, fixed in the imaging chamber, and imaged 15 minutes after Luciferin injection (150 mg/kg; i.p.). Bioluminescence signal was quantified using the Living Image software 4.2 (Caliper). 2 weeks after tumor cell injections, mice were randomly assigned to different experimental procedures, including injection of MgCl2, DNA origami conjugated with 8 Atto555 and a human carbonic anhydrase (CA) XII-specific antibody (6A10) or DNA origami conjugated with 8 Atto555 and through the tail vein, and allowed for circulation for 1 hour before perfusion.

### Whole-body clearing

**T**he labeling and clearing of CX3CR1 mice followed vDISCO protocol (6). Residual blood and heme and bone decalcification was performed by perfusion with decolorization solution and decalcification solution before staining. The decolorization solution was prepared with 25–30 vol% dilution of CUBIC reagent 15 in 0.01 M PBS. CUBIC reagent 1 was prepared with 25 wt% urea (Carl Roth, 3941.3), 25 wt% N,N,N′,N′-tetrakis (2-hydroxypropyl)ethylenediamine (Sigma-Aldrich, 122262) and15 wt% Triton X-100 in 0.01 M PBS. The decalcification solution consisted of 10 wt/vol% EDTA (Carl Roth, 1702922685) in 0.01 M PBS, adjusting the pH to 8–9 with sodium hydroxide (Sigma-Aldrich, 71687). The solutions for the labeling pipeline were pumped inside the animal’s body by transcardial-circulatory perfusion, exploiting the same entry point hole into the heart created during the PBS and PFA perfusion step (see above) and following the procedure described previoulsy6. In brief, the mouse body was placed in a 300 ml glass chamber (Omnilab, 5163279) filled with 250–300 ml of the appropriate solution, which covered the body completely. Next, the transcardial circulatory system was established with a peristaltic pump (ISMATEC, REGLO Digital MS-4/8 ISM 834; reference tubing, SC0266), keeping the pressure at 160–230 mmHg (45–60 r.p.m.). One channel from the pump, made by a single reference tube, was set for circulation of the solution through the heart into the vasculature: one ending of the tube was connected to the tip of a syringe (cut from a 1 ml syringe; Braun, 9166017V) which held the perfusion needle (Leica, 39471024), and the other ending was immersed in the solution chamber where the animal was placed. The perfusion needle pumped the appropriate solution into the mouse body, and the other ending collected the solution exiting the mouse body to recirculate the solution, pumping it back into the animal. To fix the needle tip in place and to ensure extensive perfusion, we put a drop of superglue (Pattex, PSK1C) at the level of the hole where the needle was inserted inside the heart. Using the setting explained above, after post-fixation and PBS washing, the mice were first perfused with 0.1 M PBS overnight at room temperature. Then, the animals were perfused with 250 ml of decolorization solution for 2 d at room temperature, exchanging with fresh decolorization solution every 6–12 h until the solution turned from yellowish to clear and the spleen became lighter in color (indicating that the blood heme was extracted). Then, they were perfused with 0.01 M PBS, washing for 3 h three times, followed by perfusion with 250 ml of decalcification solution for 2 d at room temperature, and again perfusion/washing with 0.01 M PBS for 3 h three times. After this, the animals were perfused with 250 ml of permeabilization solution containing 1.5% goat serum, 0.5% Triton X-100, 0.5 mM of methyl-β -cyclodextrin, 0.2% trans-1-acetyl-4-hydroxy-l-proline and 0.05% sodium azide in 0.01 M PBS for half a day at room temperature. With this setting, the animals were perfused for 6 d with 250 ml of the same permeabilization solution containing 290 μl of PI (stock concentration 1 mg ml−1) and for cancer-bearing animals, 35 μl of nanobody (stock concentration 0.5 – 1 mg/ml) to boost mCherry signal. Next, we removed the animals from the chamber, and with fine scissors, we removed a tiny piece from the back of the skull (above the cerebellum) at the level of the occipital bone. After that, the mice were washed out by perfusing with washing solution (1.5% goat serum, 0.5% Triton X-100, 0.05% of sodium azide in 0.01 M PBS) for 3 h three times at room temperature and 0.01 M PBS for 3 h three times at room temperature. After the staining, the animals were cleared using a 3DISCO-based passive whole-body clearing protocol optimized for large samples. The mice were incubated at room temperature in dehydration and clearing solutions inside a 300 ml glass chamber, kept gently rotating on top of a shaking rocker (IKA, 2D digital) inside a fume hood. For dehydration, mice bodies were incubated in 200 ml of the following gradient of THF in distilled water (12 h for each step): 50 vol% THF, 70 vol% THF, 80 vol% THF, 100 vol% THF and again 100 vol% THF, followed by 3 h in dichloromethane and finally in BABB. The glass chamber was sealed with parafilm and covered with aluminum foil during all incubation steps. Additionally, for better observation of the colocalization of CX3CR1 antibody conjugated DNA origami-Atto and CX3CR1 positive immune cells (monocytes, B cells, T cells, macrophages), liver samples were rehydrated following DCM, 100% THF, 70% THF, 50% THF and PBS x 2 times steps. The rehydrated livers were sliced with a vibratome (Leica, VT1200S) to collect several 1mm thick sections for immunostaining. These sections were blocked with blocking buffer at room temperature for 2-3 hours, a mixture of 10% goat serum, 10% DMSO (Roth, A994.2), and 0.2% Triton X-100 in PBS. 1 ml of primary antibody solution was added in a 24-well plate to the sections for 2 hours at 37 °C: rabbit antibody anti-Iba1 (Allograft inflammatory factor 1 (AIF-1))(1:1000, 019-19741, Wako) and CD86 (1:1000, Ab125212, Abcam) dissolved in the buffer of 10mg**·**L-1Heparin, 3% goat serum, 0.2%Tween-20, 3% DMSO in PBS. After three times of washing with a washing buffer containing only 0.2%Tween-20 and 10mg**·**L-1Heparin in PBS. Alexa 470-conjugated secondary antibodies (1:500, Thermo Fisher) were applied to the sample dissolved in the same buffer of primary antibody. After washing with PBS, the samples were ready for imaging. The imaging was following previous work, as summarized: “All confocal images were conducted with Zeiss LSM 880 confocal microscope mounted with a 40x oil-immersion objective (Zeiss, ECPlan-NeoFluar × 40/1.30 oil DIC M27, 1.3 NA [WD = 0.21 mm]) for higher magnification. Cleared samples were placed on the glass bottom of MatTek Petri dishes (35 mm) and immersed in several drops of BABB solution to keep the tissue transparency. All images were analyzed using Zen 2 software (v.10.0.4.910; Carl Zeiss AG).

### Light-sheet microscopy imaging

Single plane illuminated (light-sheet) image stacks were acquired using an Ultramicroscope II (LaVision BioTec), featuring an axial resolution of 4 μm with following filter sets: ex 470/40 nm, em 535/50 nm; ex 545/25 nm, em 605/70 nm; ex 560/30 nm, em 609/54 nm; ex 580/25 nm, em 625/30 nm; ex 640/40 nm, em 690/50 nm. For low magnification-whole-body imaging of the Cx3Cr1-eGFP mouse, we used a 1x Olympus air objective (Olympus MV PLAPO 1x/0.25 NA [WD = 65mm]) coupled to an Olympus MVX10 zoom body, which provided zoom-out and -in ranging from 0.63x up to 6.3x. Using 1x objective and 0.63x of zoom, we imaged a field of view of 2 × 2.5 cm, covering the entire width of the mouse body. Tile scans with 40% overlap along the longitudinal y-axis of the mouse body were obtained from ventral and dorsal surfaces up to 12 mm in depth, covering the entire body volume using a z-step of 8 μm. Exposure time was 120 ms, the laser power was adjusted depending on the intensity of the fluorescent signal (in order never to reach the saturation), and the light-sheet width was kept at a maximum. After tile imaging of the sample within the entire field of view, already scanned regions were cut using a thin motorized dental blade (0.2 mm) (Dremel 8200) for further imaging. After low magnification imaging of the whole body, lungs, liver, spleen of the MgCl2 buffer, Atto647-PEG, DNA origami-Atto647-PEG (0.5 μM) or goat anti-mouse CX3CR1 antibody conjugated DNA origami (0.5 μM) injected CX3CR1-eGFP animals were imaged with a 12x objective (Olympus MVPLAPO2XC/0.5 NA [WD = 6 mm]).

The non-targeting origami animal was scanned and imaged by mesoSPIM, an open-sourced light-sheet microscope54. The animal was scanned under 2x Zoom through Olympus MV PLAPO 1x to achieve 2x magnification. The pixel size at the object plane is 3.75 μm and the step size for z is 4 μm. The animal was mounted on a refractive-index matched glass (Ohara S-BAL14) holder to allow illumination and imaging from all different angles without retardance from the holder.

### Confocal microscopy imaging

To imaging the thick cleared specimens such as dissected tissues, pieces of organs or whole organs were placed on 35 mm glass-bottom Petri dishes (MatTek, P35G-0-14-C), the samples were then covered with one or drops of the refractive index matching solution such as BABB. Sealing of this mounting chamber was not necessary. The samples were imaged with an inverted laser-scanning confocal microscopy system (Zeiss, LSM 880) using a 40x oil immersion lens (Zeiss, EC Plan-Neofluar 40x/1.30 Oil DIC M27) and a 25x water immersion long-working distance objective lens (Leica, NA 0.95, WD = 2.5 mm), the latter one was mounted on a custom mounting thread. The z-step size was 1-2.5 mm.

### Reconstructions of full-body scans

For light-sheet microscopy reconstructions (3D montage of the entire mouse), the image stacks were acquired and saved by ImSpector (LaVision BioTec GmbH) as 16-bit grayscale TIFF images for each channel separately. The stacks were stitched with Fiji (ImageJ2) and fused with Vision4D (Arivis AG). Further image processing was done Fiji: the autofluorescence channel (imaged in 488 excitation) was equalized for a general outline of the mouse body. The organs were segmented manually by defining the regions of interest (ROIs). Data visualization was done with Amira (FEI Visualization Sciences Group), Imaris (Bitplane AG), and Vision4D in volumetric and maximum intensity projection color mapping. The images taken from mesoSPIM were stitched by Terastitcher55. Imaris was used thereafter to visualize the stitched whole-body image.

## Supporting information

Supplementary Movie 1| Single cell imaging of DNA nanotechnology in whole transparent mouse. vDISCO reveals localization of immune cell targeting DNA

Supplementary Movie 2| in vitro cellular localization of cancer-targeting DNA origami.

Supplementary Movie 3| in vivo two photon imaging of non-targeting DNA origami coated with PEG Poly(L)Lysine circulating in the mouse pial vessels.

## ACKNOWLEDGEMENTS

Illustrations were created with BioRender.com. This work was funded by the European Research Council (ERC-StGs 802305 to AL and ERC-CoG 865323 to AE), and the German Research Foundation (DFG) under Germany’s Excellence Strategy (EXC 2145 SyNergy – ID 390857198) and through FOR 2879 (ID 428662890). European Research Council Consolidator Grant to H.D. (GA no. 724261), the Deutsche Forschungsgemeinschaft through grants provided within the Gottfried-Wilhelm-Leibniz Program, and the Project ID 405686451 (to H.D.) BMBF grant HIVacToc. The contribution of NP and IK was funded by the Deutsche Forschungsgemeinschaft (DFG, German Research Foundation) under Germany’s Excellence Strategy within the framework of the Munich Cluster for Systems Neurology (EXC 2145 SyNergy – ID 390857198). M.M. is supported by Turkish Ministry of Education for her PhD studies.

## AUTHOR CONTRIBUTION

M.M. performed and analyzed most of the experiments. B.K. designed and produced all DNA origami particles. S.Z., V.S., and H.S.B. performed sample collection and experiments in the project’s initial phase. S.Z. Performed IHC of liver samples. T-L.O. acquired whole-body images of non-targeted DNA origami and reconstructed with M.I.T. S.R., A.S., and V. S. performed flow cytometry for immunogenicity assessments. I.K. performed 2-photon intravital imaging of DNA origami in pial vessels. C.P. S.Z. and M.M. performed the transcardiac injection of MDA-MB-231 cells. R.Z. provided the 6A10 antibody. F.H. supervised part of the project and helped write the manuscript. A.L. designed and supervised part of the experiments (flow cytometry, FACS) and edited the manuscript. NP designed part of the experiments (2-PM) and edited the manuscript. F.H. A.E., H.D., and M.M. co-wrote the final manuscript. A.E. and H.D. designed and led all aspects of the research. All authors read and approved the final manuscript.

## DECLARATION OF INTERESTS

Authors declare no competing interests.

## MOVIE LEGENDS

**Supplementary Movie 1**| Single cell imaging of DNA nanotechnology in whole transparent mouse. vDISCO reveals localization of immune cell targeting DNA origami in 3D reconstruction of whole transparent mouse body. Muscles are labeled in cyan, bones and organs in white and DNA origami in red. DNA Origami conjugated with anti-mouse CX3CR1 antibody is mostly localized in immune cells in the spleen, the liver and the lungs. Similar results were observed in 2 independent animals.

**Supplementary Movie 2**| in vitro cellular localization of cancer-targeting DNA origami. MDA-MB-231 cells cultured with cancer-targeting DNA origami and time lapse imaging 24 hours using confocal microscope. MDA-MB-231 mCherry cells are in cyan and cancer-targeting DNA origami is in red. DNA origami binds the surface of the cells and then internalized.

**Supplementary Movie 3**| in vivo two photon imaging of non-targeting DNA origami coated with PEG Poly(L) Lysine circulating in the mouse pial vessels. Meningeal vessels recorded through acute cranial window for 70 minutes (green -FITC-dextran) before and after injection of DNA origami Atto 647N; 80 μm depth. Laser power 3.5%-10%.

**Figure S1.**
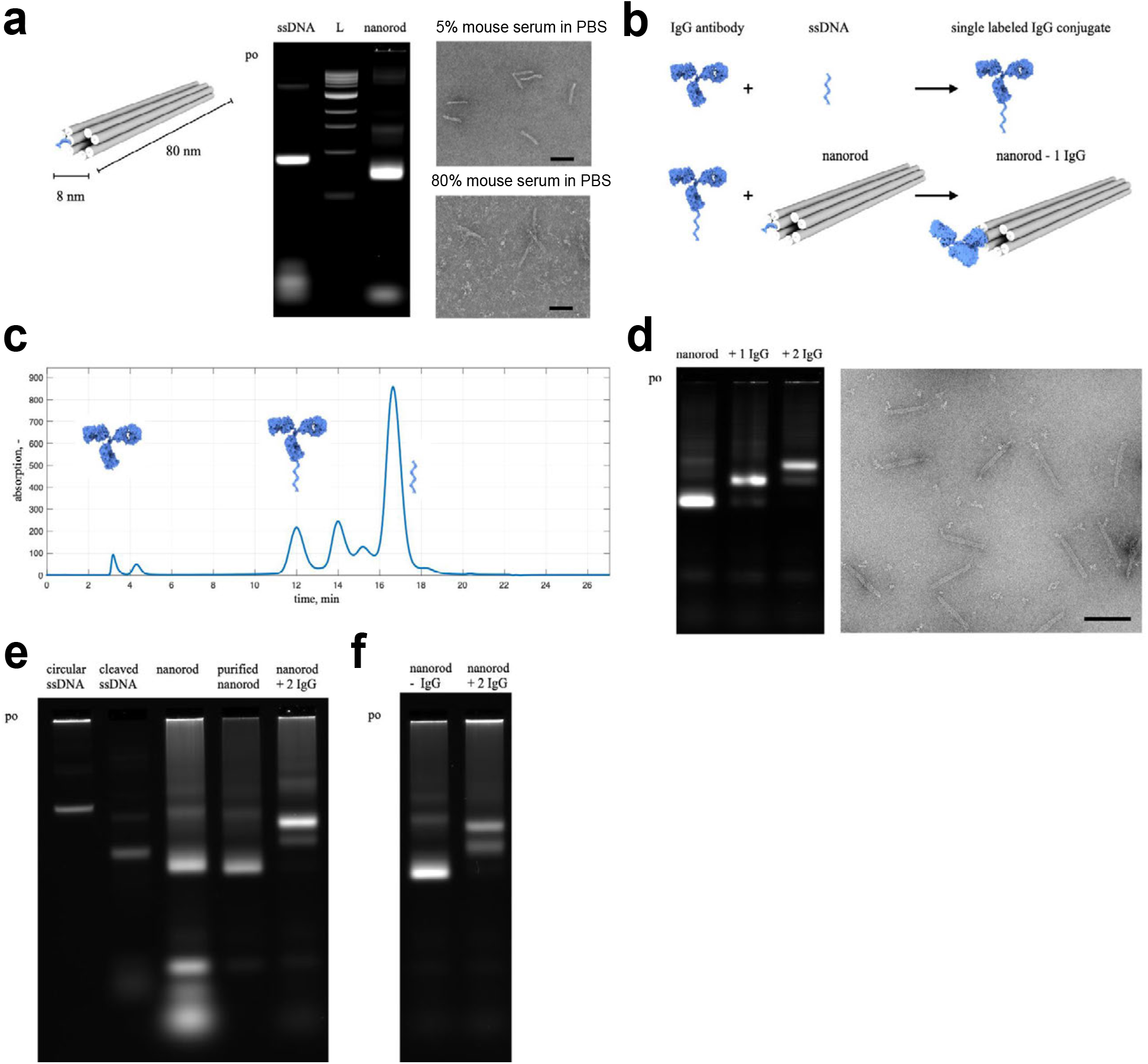
Antibody conjugation and production of DNA origami nanorod for *in vivo* studies. **a**, left: Schematic representation of DNA origami nanorod used in *in vivo* studies. middle: Laser scanned image of 2 % agarose gel with from left to right the cleaved phagemid ssDNA yielding scaffold and staple strands, a 1kb ladder as reference and the folded nanorod. right: negative stain transmission electron microscope (TEM) micrograph of the nano rod stabilized with PEG-polylysine and incubated in mouse serum (5% (top), 80% (bottom), at 37°C, 1h, 50 fold diluted for grid preparation) (scale bar 100 nm). **b**, Schematic representation of attaching single stranded DNA to IgG antibody and hybridization to nanorod using single stranded adapter protruding from a helix end. **c**, Purification of the coupling reaction of IgG antibody and single-stranded ssDNA via anion exchange chromatography. The peak between 2.5 and 5 min in the chromatogram corresponds to the unmodified antibody, the peak at 12 min to the antibody modified with one ssDNA strand, which represent the product of the coupling reaction. At 14 min the antibody modified with 2 DNA strands appears as a byproduct. The peak at 17 min corresponds to the unreacted DNA present in excess. Single and double labeled antibody was verified using SDS-PAGE and agarose gel analysis (data not shown). **d**, left: Agarose gel on which the nanorod with 0 to 2 protruding single stranded handle adapters, incubated with IgG modified with one ssDNA strand (1.5 x excess over binding site), was electrophoresed (po, pocket). right: TEM micrograph of nanorod with one IgG antibody attached (scale bar 100 nm). **e**, Laser scanned image of a 2% agarose gel on which from left to right the biotechnological produced circular phagemid ssDNA, the cleaved phagemid ssDNA, the folded nanorod with additional chemically synthesized fluorescent oligos (Atto-550), the PEG purified nanorod and the nanorod with 2 binding sites for IgG antibody 6A10 was analyzed. **f**, Exemplary quality control of the PEG purified nanorod without and with two 6A10 antibodies attached for an *in vivo* experiment.

**Figure S2.**
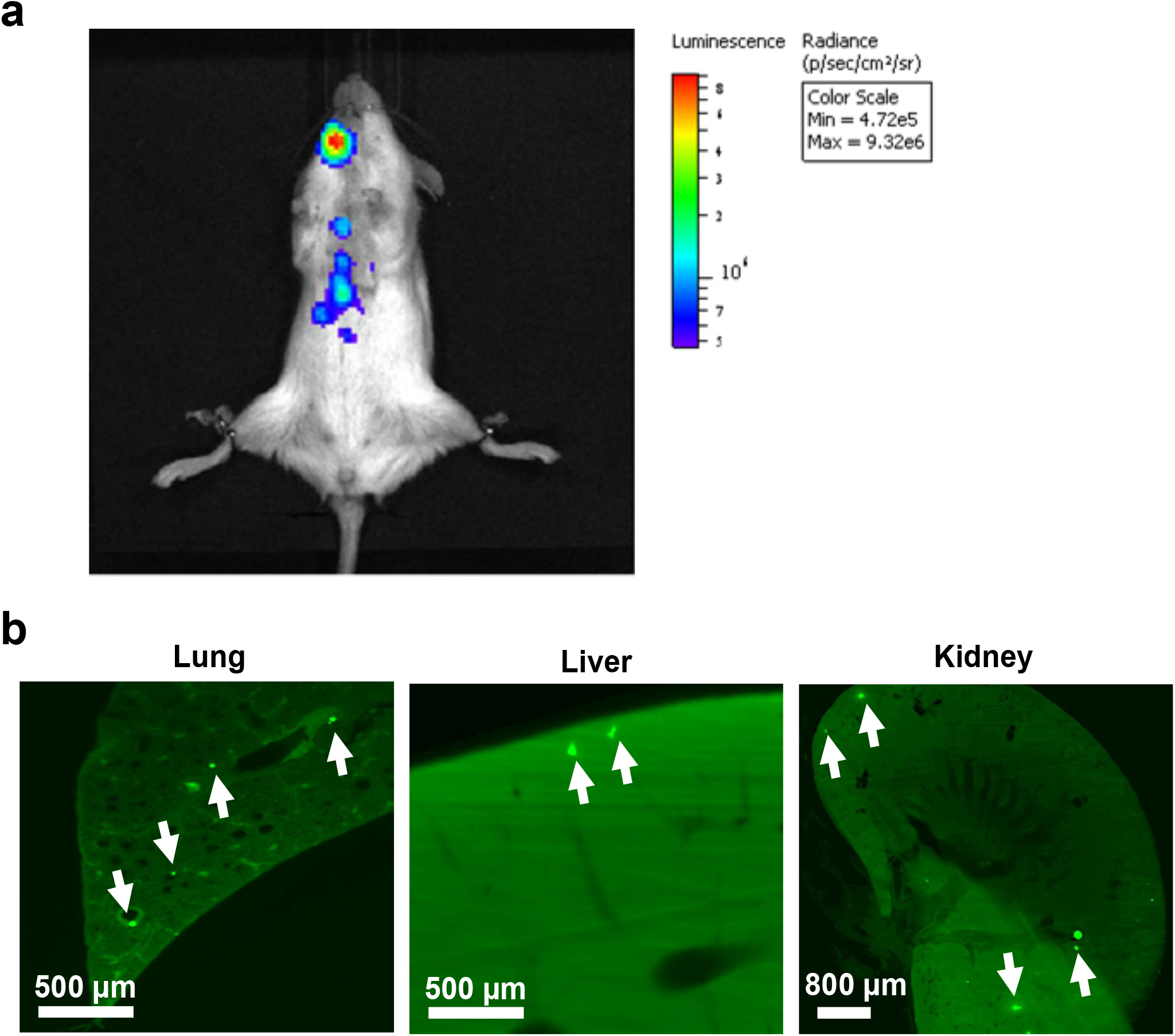
Visualization of metastases in a full-body. **a**, *in vivo* bioluminescence imaging of a female NSG mouse 2 weeks after MDA-MB-231 cancer cell implantation through transcardiac route. **b**, vDISCO enable to identify single micrometastases in the same mouse in internal organs such as lung, liver and kidney. MDA-MB-231 cells are green.

**Figure S3.**
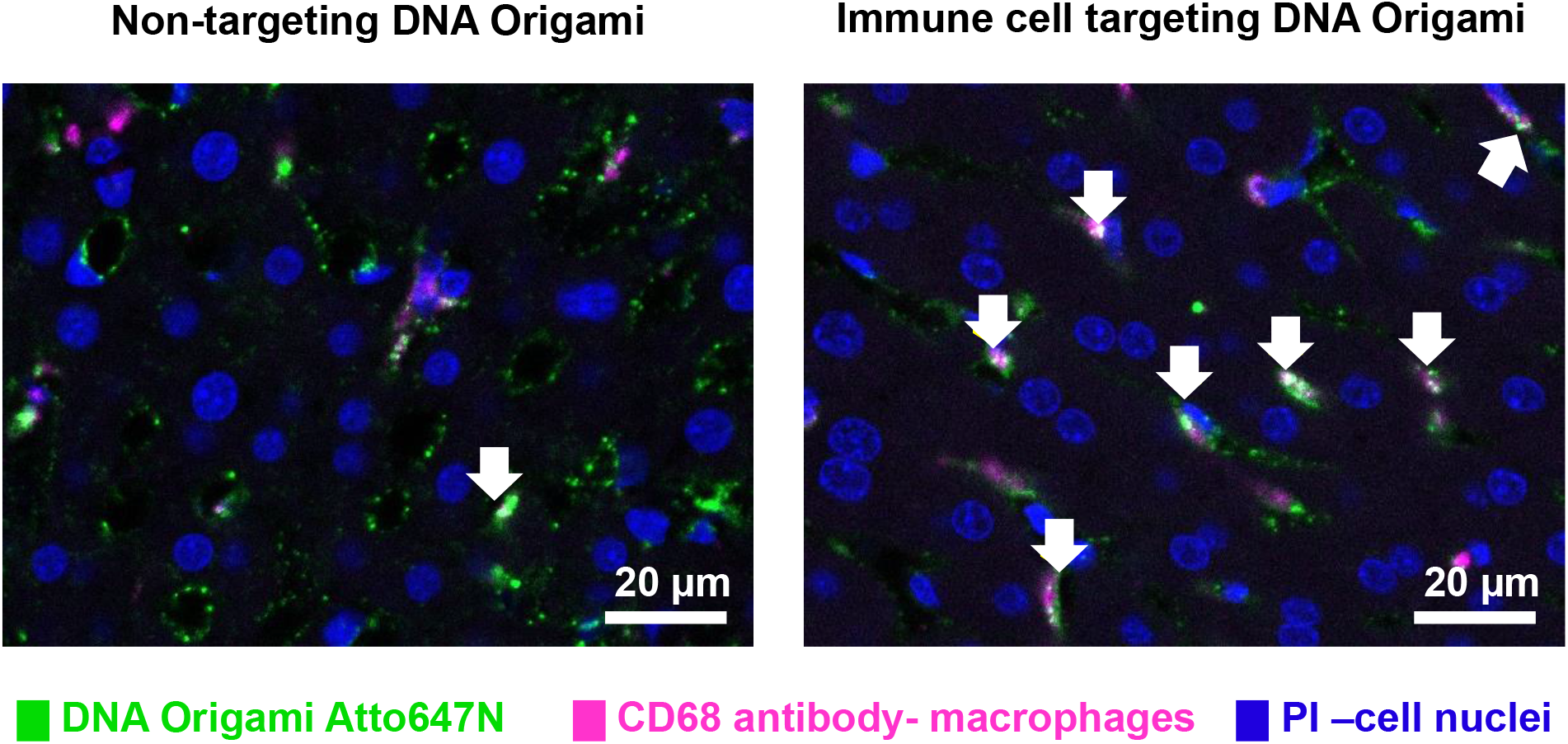
Addressability of DNA Origami. CX3CR1 positive tissue macrophages are evaluated by immunohistochemistry in 1mm sections of livers with the macrophage lysosomal vesicle specific CD68 antibody in non-targeting DNA Origami Atto647 and immune cell targeting Origami injected animals. Engulfment of DNA Origami CX3CR1 antibody Atto647 PEG-Polylysine with higher efficiency can be observable with colocalization of DNA Origami signal and CD68 labeling (arrows).

**Figure S4.**
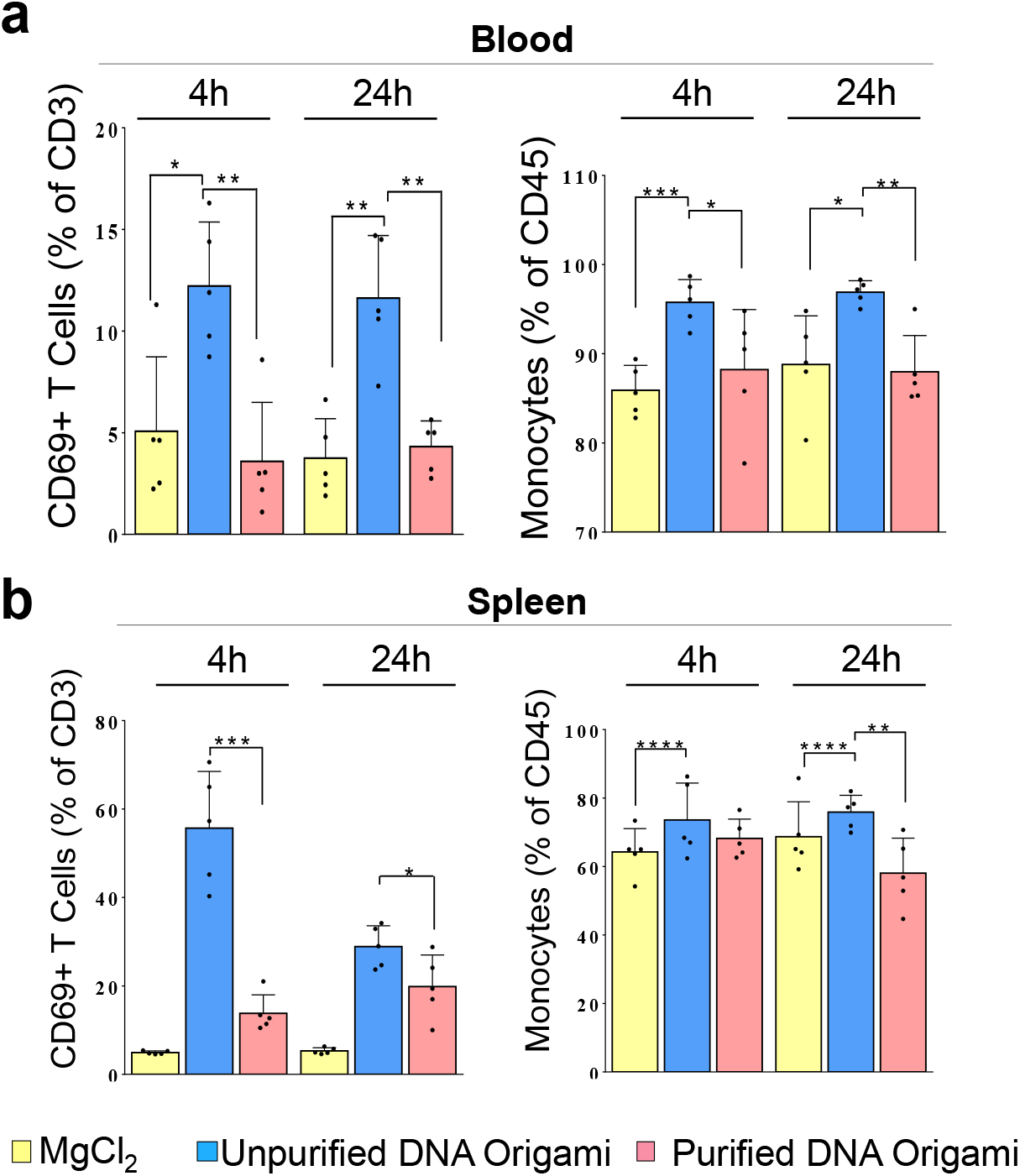
Assessment of immune safety of DNA Origami. Comparison of the immune reaction against the MgCl2 (folding solution), unpurified Origami containing endotoxins from E.coli culture and purified Origami. Comparison of monocytes and activated T cell counts are represented for a, blood and b, spleen in at acute stage after DNA Origami administration through tail-vein injection (n=5). Purification efficiently resolve Origami to be immune tolerable. (*: p ≤ 0.05, **: p ≤ 0.01, ***:p ≤ 0.001, ****:p ≤ 0.0001, following one-way ANOVA)

**Figure S5.**
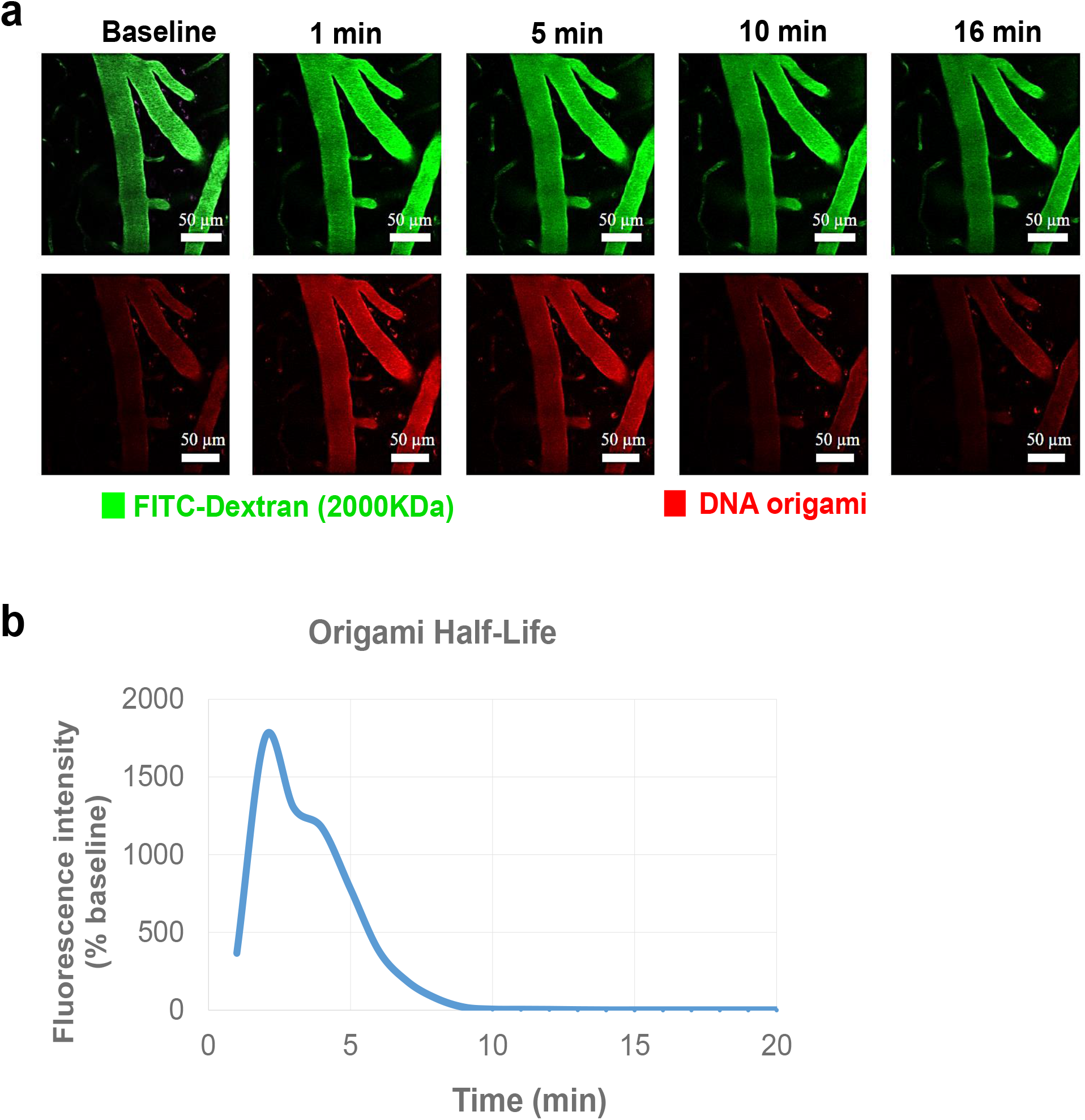
Half-life of DNA origami in blood. **a**, Blood clearance of non-targeting DNA Origami by intravital 2-photon imaging of the meningeal vessels after intravenous injection of FITC Dextran (2000 KDa) to label the vessels and Naked DNA Origami. After the injection of non-targeting origami, the fluorescent signal was recorded during the next 70 minutes. The origami signal immediately increased upon injection and then returned to baseline level within 16 minutes. **b**, Quantification of relative intravascular fluorescence intensities over time showed that the intensity of DNA Origami-Atto647 steeply decreased 5 minutes after injection and further becoming negligible after 10 minutes.

## Notes

### Competing Interest Statement

The authors have declared no competing interest.

